# Pre-clinical proof-of-concept of anti-fibrotic activity of caveolin-1 scaffolding domain peptide LTI-03 in *ex vivo* precision cut lung slices from patients with Idiopathic Pulmonary Fibrosis

**DOI:** 10.1101/2024.04.24.589970

**Authors:** BreAnne MacKenzie, Poornima Mahavadi, Yago Amigo Pinho Jannini-Sa, Brecht Creyns, Ana Lucia Coelho, Milena Espindola, Clemens Ruppert, Konrad Hötzenecker, Cory Hogaboam, Andreas Guenther

## Abstract

**Rationale:** While rodent lung fibrosis models are routinely used to evaluate novel antifibrotics, these models have largely failed to predict clinical efficacy of novel drug candidates for Idiopathic Pulmonary Fibrosis (IPF). Moreover, single target therapeutic strategies for IPF have failed and current multi-target standard of care drugs are not curative. Caveolin-1 (CAV-1) is an integral membrane protein, which, via its caveolin scaffolding domain (CSD), interacts with caveolin binding domains (CBD). CAV-1 regulates homeostasis, and its expression is decreased in IPF lungs. LTI-03 is a seven amino acid peptide derived from the CSD and formulated for dry powder inhalation; it was well tolerated in normal volunteers (NCT04233814) and a safety trial is underway in IPF patients (NCT05954988). **Objectives:** Anti-fibrotic efficacy of LTI-03 and other CSD peptides has been observed in IPF lung monocultures, and rodent pulmonary, dermal, and heart fibrosis models. This study aimed to characterize progressive fibrotic activity in IPF PCLS explants and to evaluate the antifibrotic effects of LTI-03 and nintedanib in this model. **Methods:** First, CBD regions were identified in IPF signaling proteins using *in silico* analysis. Then, IPF PCLS (n=8) were characterized by COL1A1 immunostaining, multiplex immunoassays, and bulk RNA sequencing following treatment every 12hrs with LTI-03 at 0.5, 3.0, or 10 μM; nintedanib at 0.1 μM or 1 μM; or control peptide (CP) at 10 μM. **Measurements and Main Results:** CBDs were present in proteins implicated in IPF, including VEGFR, FGFR and PDGFR. Increased expression of profibrotic mediators indicated active fibrotic activity in IPF PCLS over five days. LTI-03 dose dependently decreased COL1A1 staining, and like nintedanib, decreased profibrotic proteins and transcripts. Unlike nintedanib, LTI-03 did not induce cellular necrosis signals. **Conclusion:** IPF PCLS explants demonstrate molecular activity indicative of fibrosis during 5 days in culture and LTI-03 broadly attenuated pro-fibrotic proteins and pathways, further supporting the potential therapeutic effectiveness of LTI-03 for IPF.

## Introduction

Idiopathic pulmonary fibrosis (IPF) is an interstitial lung disease (ILD) that is estimated to affect over 200,000 people in the United States and 1 out of every 200 adults over the age of 60 (1). IPF is characterized by progressive interstitial fibrosis of the lung parenchyma which results in continuous loss of lung function and gas exchange properties. As a result, patients experience dry coughing, progressive dyspnea, initially under exercise, later at rest, and, ultimately, death due to respiratory failure. In untreated patients, the median survival from the time of diagnosis is 2-3 years (1). Hence, there is a great unmet medical need.

Following the identification of a potential candidate therapeutic for IPF, drugs are typically evaluated for antifibrotic efficacy in the mouse bleomycin (BLM) model of lung fibrosis. A statistically significant reduction in mouse lung hydroxyproline content following therapeutic intervention has been a prerequisite for investment in further clinical development. Although the BLM model is useful for determining whether a compound has antifibrotic activity in vivo, it is not predictive of a successful treatment outcome for IPF and this is due to several reasons: **i)** IPF is a prolonged, complex disease which occurs over decades in an aging lung often in the context of exposure to multiple environmental insults and genetic insufficiencies; the BLM model is typically performed from 7 to 28 days in young, female mice of an inbred genetic background, housed in clean and sterile facilities, **ii)** there are striking differences with regard to disease onset (drug toxicity in the BLM model; endoplasmic, lysosomal or DNA damage in IPF) and **iii)** the natural course of disease (BLM model resolves spontaneously while IPF is characterized by a progressive decline in lung function due to unresolved scarring). While more than 700 drugs have demonstrated efficacy in the BLM model, and over 145 clinical trials for IPF have taken place, only two drugs, Pirfenidone (Esbriet®) (1, 2) and Nintedanib (OFEV®) (3), have been authorized. While both drugs slow down disease progression and expand survival time, they fail to entirely stop disease progression or to induce lung regeneration. Therefore, the development and characterization of a translational model of IPF that is truly predictive of clinical efficacy would be highly beneficial.

Caveolin-1 (CAV1) is an integral membrane protein expressed primarily in epithelial, endothelial, immune cells and stromal cells of the lung. (8) CAV1 is required for the formation of plasma membrane invaginations (caveolae) (9) and is implicated in receptor endocytosis, signaling cascades, membrane transport and immunity. Loss of CAV-1 mRNA and protein expression has been observed in many cell types of human IPF lung tissue (4) as well as in many other human tissues affected by fibrotic disease (5). In addition, Cav-1 expression is reduced in a rodent bleomycin model of pulmonary fibrosis (6) and *Cav1^-/-^* mice develop spontaneous pulmonary fibrosis (PF) (7).

A hydrophobic, seven amino acid sequence (FTTFTVT), referred to as CSP7 or LTI-03, has been derived from the broad interacting caveolin scaffolding domain (CSD), which putatively binds to any endogenous proteins predicted to have CSD binding domains (CBD.) (10). CBDs play a role in CAV-1 dimerization as well as regulation of diverse signaling intermediates, many of which are implicated in the pathogenesis of fibrosis (4). LTI-03 was designed to replenish the modulating, homeostatic effects of the CSD domain in the IPF lung. Indeed, LTI-03 was identified based on the inhibitory effect of CSD peptide truncations, deletions, and substitutions on ECM production by human IPF and bleomycin-injured mouse lung fibroblasts and retained the anti-fibrotic activity of full length CSD. (11) Moreover, in rodent cardiac, pulmonary, and dermal (12–14) fibrosis models, systemic or local administration of LTI-03 prevented or attenuated organ fibrosis. LTI-03 was formulated as an excipient-free dry powder for inhalation for the treatment of IPF (15). It mitigated pulmonary fibrosis (PF) in both chronic and acute mouse models (16) and has demonstrated anti-fibrotic effect in IPF epithelial and fibroblast monocultures (17, 18) but its activity had yet not been assessed in intact IPF lung tissue.

In this report, we used human *ex vivo* IPF precision cut lung slices (PCLS) to rigorously test anti-fibrotic activity and to analyze the downstream signaling machinery of LTI-03 in a novel and innovative 3D culture of viable IPF tissue. Progressive fibrotic signaling was observed in untreated human IPF PCLS over five days in culture. IPF PCLS treated with either LTI-03 or the standard of care nintedanib, demonstrated comparable anti-fibrotic activity as indicated by diminished COL1A1 immunofluorescence and attenuation of proinflammatory and profibrotic transcript and protein expression. LTI-03 had similar pleiotropic antifibrotic effects compared to the standard of care (SOC) treatment, nintedanib, without any evidence of cellular necrosis or toxicity. Taken together, we hypothesize that use of human *ex vivo* IPF PCLS represents an innovative translational assay to evaluate proof of concept of drug candidates and may, in the future, serve as a preclinical, predictive model of clinical efficacy.

## Methods

### In silico identification of caveolin-1 binding domains in human proteome

CBD motif 1 (φXφXXXXφ), CBD motif 2 (φXXXXφXXφ), and the CBD composite motif (φXφXXXXφXXφ) were identified using the MOTIF Search tool (setting: nr-aa) available on GenomeNet (genome.jp) (19). The symbol ‘φ’ represents either Tryptophan (Trp - W), Phenylalanine (Phe – F), or Tyrosine (Tyr – Y), while X is any other amino acid. These sequences were used as queries to identify protein sequences containing CBDs (20).

### Precision cut lung slice (PCLS) cultures

Human lung tissues from idiopathic pulmonary fibrosis (IPF) patients undergoing lung transplantation and non-IPF donors were obtained via the European IPF Registry (euIPFreg) sites Vienna and Giessen and delivered via the European IPF Biobank or the UGMLC Biobank, a member of the DZL platform biobanking. The study protocol was approved by the Ethics Committee of the Justus-Liebig University School of Medicine (No. 111/08: European IPF Registry and 58/15: UGMLC Giessen Biobank), and informed consent was obtained in written form from each subject (a comprehensive list of patient data and applied assays is provided in **Table 2**). Lung biopsy cores were obtained exclusively from 8 patients undergoing lung transplant for treatment of IPF and processed for PCLS culture as previously described (21). All patients were male, an average of 61.5 years of age (stdev 6.56 years), and had a clinical diagnosis of IPF alone, or IPF with secondary pulmonary hypertension (PH). Usual interstitial pneumonia (UIP) was the main pathological finding on all explant tissue (**Table 2**). The models and assays performed in each PCLS culture are also summarized in **Table 2** (22).

### PCLS treatments

LTI-03 (Sequence: FTTFTVT; Lot# AJF73B//474-33-TI344 Lonza), the inactive control peptide CP1 (Sequence: FAAFAVA; Lot# PD200120-1, Ambiopharm) were suspended in PCLS culture media at 0.3, 5.0, or 10μM for treatments. Nintedanib (Boehringer Ingelheim Pharmaceuticals, Inc #S1010) was prepared in a 1mM stock solution with sterile filtered DMSO and used at 0.1 μM or 1 μM. PCLS were treated every 12hrs and supernatant was harvested 12hrs following the last of either three (day 2), seven (day 4) or nine treatments (day 5). Tissue was harvested after 9 treatments (day 5).

### Collagen 1 alpha 1 immunofluorescence quantification

Collagen 1 alpha 1 (COL1A1) immunostaining was performed on tissue sections generated from 4 PCLS biological samples. Immunofluorescence using a COL1A1 (Rockland) monoclonal antibody was performed as described (22). Leica M205 FA Fluorescent Stereoscope (Leica Microsystems) and LAS-X-Core Software (3.7.4. version, LAS X Life Science, Leica Microsystems) was used for imaging. Five pictures from each stained PCLS were obtained and quantified with ImageJ on converted 16-bit images. Five regions from each image were randomly selected and measured and mean values were calculated. At least 5 technical replicates were used per treatment group. Statistics indicate were reported as the standard error of the mean.

### Bio-plex

Proteomic analysis on 8 unique IPF PCLS supernatants collected at days 2, 4 or 5 after treatment was performed using Luminex bead-based assay kit (Bio-Plex Pro™ Human Inflammation Panel 1, #171AL001M, Bio Rad) according to manufacturer’s instructions. Data was acquired using a Bio-Plex 200 reader (Bio Rad) and are presented as the ratio compared to untreated (control) PCLS.

### Bulk-RNA-sequencing

Bulk RNA sequencing was performed on tissue from 4 separate PCLS. Briefly, PCLS were minced manually in 2 mm pieces in Trizol (ThermoFisherScientific) before bead homogenization for 95 seconds (40+20+15+10+10 seconds) at 50 Hz. RNA was purified using a column-based workflow (Machery Nagel, 740984.250). RNA quantity and quality was assessed by RNA integrity number (RIN) via 4200 TapeStation system (Agilent). RNAseq libraries were prepared using Truseq stranded mRNA kit (Illumina; 20020594). Sequencing libraries were pooled and submitted for next-generation sequencing of 100 bp single-end reads (Illumina NovaSeq sequencer, AGCT Core; Los Angeles, CA).

### Ingenuity Pathway Analysis (IPA)

Files containing differentially expressed genes (DEGs) were generated comparing each treatment to its respective untreated group (UT) using DESeq2 package. Files were uploaded to Ingenuity Pathway Analysis (IPA; QIAGEN Inc.) for further analysis. Venn diagrams were generated comparing a dataset of Treatment vs UT against the respective day of Nintedanib or LTI-03 versus UT. Canonical pathways, Upstream Regulators and Disease & Function analysis was made using nintedanib or LTI-03 versus UT as reference for z-score and –log(p-value). Any feature with a –log(p-value) higher than 1.3 and a z-score higher than 2 or lower than −2 is significant.

### Statistical methods

Semi-quantitative results were graphed using GraphPad Prism 9.0 Software. Where appropriate a Bonferroni post-hoc analysis was performed.

## Results

### *In silico* identification of putative LTI-03 targets in IPF

LTI-03 is a seven amino acid peptide derived from the CSD region of caveolin-1, which putatively interacts with proteins containing complimentary CBD (10). To elucidate the proteins LTI-03 may be interacting with, CBD sites were queried using the MOTIF Search tool (setting: nr-aa) available on GenomeNet (genome.jp) (19) (21). A total of 7,634 proteins in the human proteome were predicted to contain these CBD sequences (**Supplemental Table 1**). Multiple CBD domains were found in the following proteins of relevance to the pathogenesis of IPF: fibroblast growth factors (FGFs), fibroblast growth factor receptors 1-4 (FGFR1-4), frizzled (FZD; WNT receptors), platelet derived growth factor receptors A and B (PDGFR), and vascular growth factor receptors 1 and 3 (VEGF). FGFR1-4, PDGFA/B, VEGFR1/3 are known targets of Nintedanib (23) and contain CBDs as shown in **Table 1**.

**Table 1.** CBD binding motifs relevant for IPF and indication of known targets of nintedanib.

| GENE | UNIPROT | CBD motif 1<br>$\phi X \phi XXXX \phi$ | CBD motif 2<br>$\phi XXXX \phi XX \phi$ | mixed domain<br>$\phi X \phi XXXX \phi XX \phi$ | Nintedanib<br>targets |
| --- | --- | --- | --- | --- | --- |
| FGF2 | P09038 | 0 | 1 | 0 |  |
| FGF5 | P12034 | 0 | 1 | 0 |  |
| FGF8 | P55075 | 0 | 1 | 0 |  |
| FGF9 | P31371 | 0 | 1 | 0 |  |
| FGF11 | Q92914 | 0 | 1 | 0 |  |
| FGF12 | P61328 | 1 | 1 | 0 |  |
| FGF13 | Q92913 | 1 | 1 | 0 |  |
| FGF14 | Q92915 | 1 | 1 | 0 |  |
| FGF16 | O43320 | 0 | 1 | 0 |  |
| FGF20 | Q9NP95 | 0 | 4 | 0 |  |
| FGFR1 | P11362 | 2 | 1 | 1 | • |
| FGFR2 | P21802 | 1 | 1 | 1 | • |
| FGFR3 | P22607 | 1 | 1 | 1 | • |
| FGFR4 | P22455 | 1 | 1 | 1 | • |
| FZD1 | Q9UP38 | 1 | 0 | 0 |  |
| FZD2 | Q14332 | 1 | 0 | 0 |  |
| FZD4 | Q9ULV1 | 1 | 2 | 1 |  |
| FZD5 | O75084 | 1 | 0 | 0 |  |
| FZD6 | Q9H461 | 3 | 0 | 0 |  |
| FZD7 | O00144 | 1 | 2 | 1 |  |
| FZD10 | Q9ULW2 | 0 | 1 | 0 |  |
| PGFRA | P16234 | 1 | 3 | 1 | • |
| PGFRB | P09619 | 1 | 2 | 1 | • |
| VGFR1 | P17948 | 3 | 1 | 1 | • |
| VGFR3 | P35916 | 2 | 2 | 1 | • |

**Table 2.** Key patient data and applied models and assays.

| key patient data |  |  |  | applied models and assays |  |  |  |
| --- | --- | --- | --- | --- | --- | --- | --- |
| age | sex | clinical diagnosis | pathological fibrosis pattern in explant tissue | PCLS | COL1α1 immuno-fluoresence | Bio-Plex Human Inflammation Panel 1 | bulk RNA sequencing |
| 53 | M | IPF | UIP | • |  | • | • |
| 58 | M | IPF | UIP | • |  | • | • |
| 60 | M | IPF, PH | UIP | • |  | • | • |
| 70 | M | IPF | UIP | • |  | • | • |
| 70 | M | IPF, PH | UIP | • | • | • |  |
| 57 | M | IPF, PH | UIP | • | • | • |  |
| 66 | M | IPF | UIP | • | • | • |  |
| 56 | M | IPF | UIP | • | • | • |  |

### Profibrotic proteins and transcripts increased in a temporal manner in IPF PCLS cultures

To determine whether IPF PCLS exhibited dynamic changes in profibrotic and proinflammatory mediators over the course of the culture time examined in this study, a Bio-Plex Human Inflammation Panel I was used to measure mediators in supernatants at days 2 and 5 of PCLS culture. Most inflammatory and pro-fibrotic proteins were increased on day 5 compared to day 2, suggestive of a progressive profibrotic activity in PCLS cultures over this time frame (**Figure 1A**). In addition, transcript analysis via bulk RNA-sequencing revealed that factors relevant to ECM deposition, secreted factors, and intracellular and plasma membrane factors related to fibrosis, were also increased over the time of PCLS culture. (**Figure 1B**). Profibrotic factors containing CBD domains were also increased (**Figure 1C**). These data provide evidence that the PCLS system adequately reflects the progressive nature of the human disease in the explanted PCLS in the culture dish.

**Figure 1.**
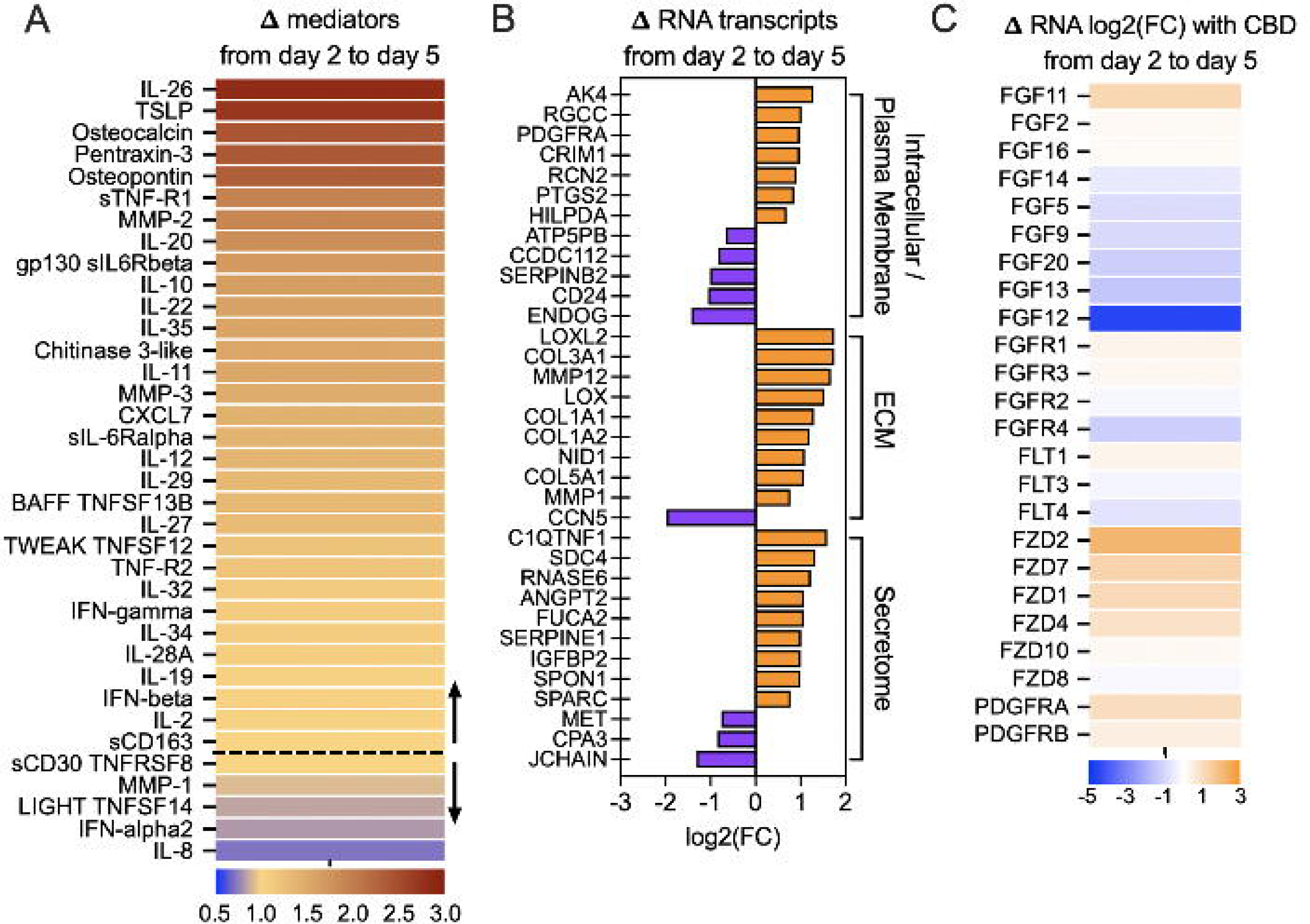
Progressive fibrotic activity was observed in IPF PCLS cultures over 5 days. *(A)* Ratio of the change in the expression of proinflammatory and profibrotic mediators from day 2 to day 5 in supernatant harvested from untreated, IPF PCLS explants (n=8 unique biological samples). (*B*) Log2 fold change (FC) of gene expression changes related to ECM deposition, secreted factors and intracellular and plasma membrane factors were increased from day 2 to day 5 according to bulk RNA sequencing of IPF PCLS explants (n=4 unique biological samples). (*C*) Profibrotic factors containing CBD domains were also increased.

### LTI-03 dose dependent reduction of COL1A1 protein in IPF PCLS tissue following 5 days in culture

Diseased, fibrotic lung fibroblasts present in the IPF lung deposit COL1A1, an extracellular matrix protein which destroys lung architecture and leads to a decline in lung function (24). In IPF PCLS (n=4), CP (at 10 μM), LTI-03 (at doses of 0.5, 3, or 10 μM) or nintedanib (0.1 μM or 1 μM) was administered to IPF PCLS every 12 hours for 5 days and representative images from two IPF patient PCLS studies (IPF1029 and IPF1030) are shown in **Figure 2A-N**. At doses of 3 (**Figure 2F & M**) or 10μM (**Figure 2G & N**), LTI-03 treatments visibly decreased COL1A1 staining intensity compared to the untreated (**Figure 2A & H**) and CP-treated (**Figure 2D & K**) groups. Staining intensity for COL1A1 also appeared to be decreased in the 0.1 μM (**Figure 2B & I**) and 1 μM (**Figure 2C & J**) nintedanib treatment groups. Calculations of fluorescence intensity were performed on a minimum of 5 areas per PCLS section and 3 PCLS technical replicates per treatment group, and these data are summarized in **Figure 2O**. Both, the 3 and 10 μM LTI-03 treatments significantly decreased COL1A1 staining intensity compared with the untreated groups and with CP (**Figure 2O**).

**Figure 2.**
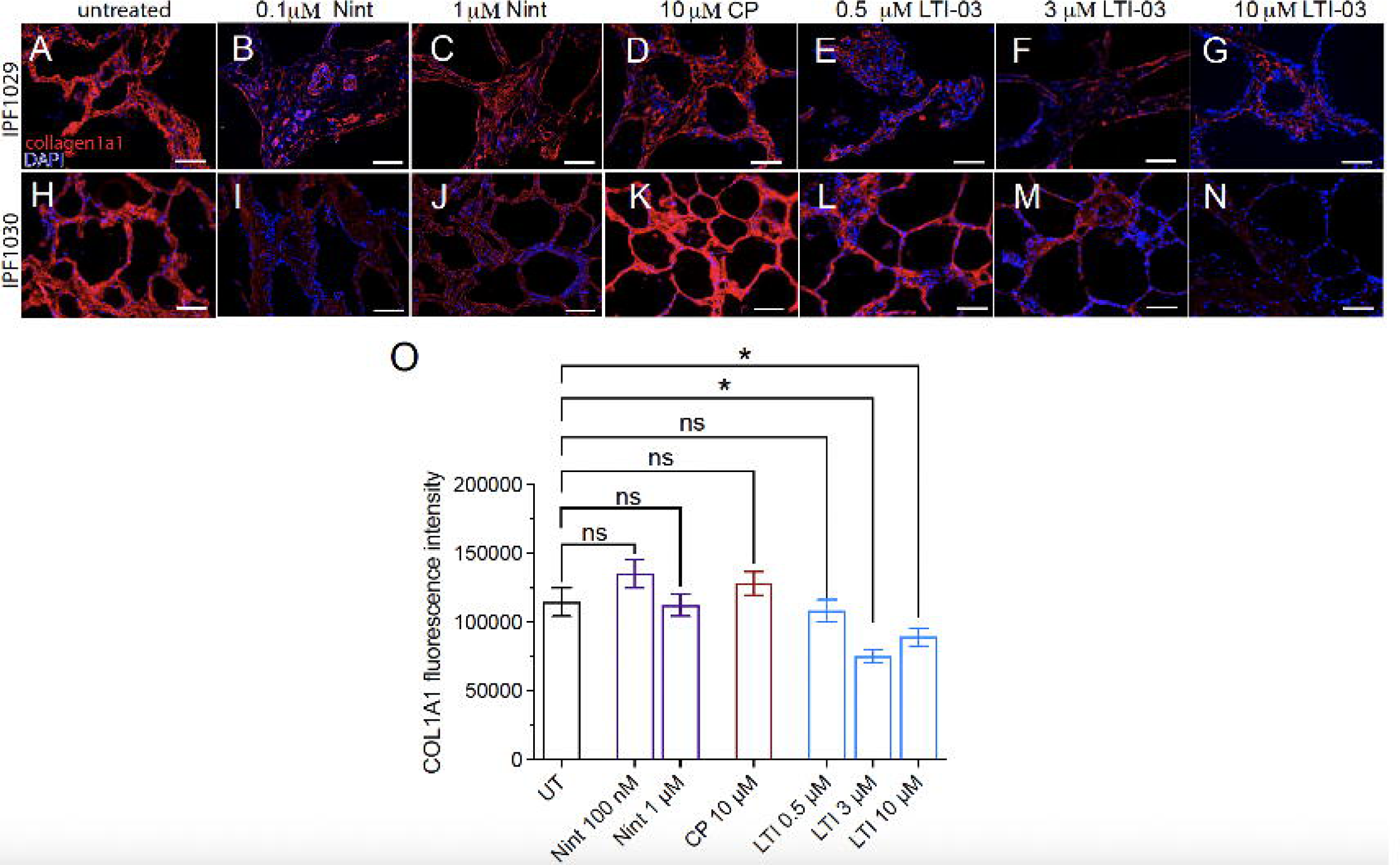
Dose dependent decrease of Collagen1α1 protein in LTI-03 treated IPF PCLS after five days in culture. Representative immunostaining staining of 2 IPF PCLS for collagen1α1 (red) with nuclear DAPI counterstain (blue) and all treatment groups: (*A, H*); untreated, (*B, I*); 0.5 μM LTI-03; (*C,J*); 3.0 μM LTI-03, (*D,K*); 10.0 µM LTI-03; (*E,L*); 10.0 μM CP; (*F,M*); 0.1 μM Nintedanib; 1.0 μM Nintedanib; Scale bars 400 μm. (*O*) Quantification of fluorescence intensity \**P*,<0.01. Error bars represent standard error of the mean.

### LTI-03 inhibits profibrotic and inflammatory mediators in IPF PCLS supernatants after 2 and 5 days of treatment

Cell free supernatant was removed from PCLS tissue cultures on day 2 and 5 post treatment and quantified for inflammatory and profibrotic mediators using a human Bio-plex panel. The heatmap indicates the mean values per protein measured normalized to the mean of untreated controls. As shown in **Figure 1**, Nintedanib had the strongest, most broad acting, dose-dependent, suppressive effect on inflammatory and profibrotic mediators after 5 days and this was followed by LTI-03 at a dose of 10μM. In contrast, CP increased levels of many of these factors. Similar results with all treatments and controls were observed following day 2 measurements (**Supplemental Figure 1**).

### Canonical pathways modulated in IPF PCLS cultures following Nintedanib or LTI-03 treatments

Ingenuity pathway analyses (IPA) of RNAseq datasets revealed that various canonical signaling pathways were modulated in IPF PCLS tissue after 5 days of nintedanib (0.1 or 1.0 μM), control peptide (10.0 μM), or LTI-03 (0.4, 3, or 10 μM) treatments every 12 hrs (n=4/group)(**Figure 4**). Treatment suppressed IPA pathways are summarized in **Figure 4A**, and treatment activated IPA pathways in **Figure 4B**. All comparisons were calculated between treated and untreated groups. LTI-03 dose-dependently inhibited the Idiopathic Pulmonary Fibrosis Signaling Pathway, which is comprised of transcripts such as TGFβ, VEGFA, PDGFB, EGF, IL6, IL1β, and FGF2 (**Figure 4A**). Other key fibrosis pathways that were consistently inhibited by LTI-03 included Pulmonary Healing Signaling Pathway, Osteoarthritis Pathway, Ephrin Receptor Signaling, Leukocyte Extravasation Signaling, FcγR-mediated Phagocytosis, and Regulation of Actin-based Motility by Rho (**Figure 4A**). LTI-03 also upregulated canonical pathways particularly PTEN, PPAR, and Xenobiotic Metabolic PXR signaling pathways, albeit these effects were mostly apparent at the 3μM dose. Compared with LTI-03 treatments, Nintedanib at a dose of 1μM exhibited stronger suppression or activation of the canonical pathways summarized in **Figure 4**. Day 2 analysis summarizing these canonical pathways is available in **Supplementary Figure 2**.

**Figure 3.**
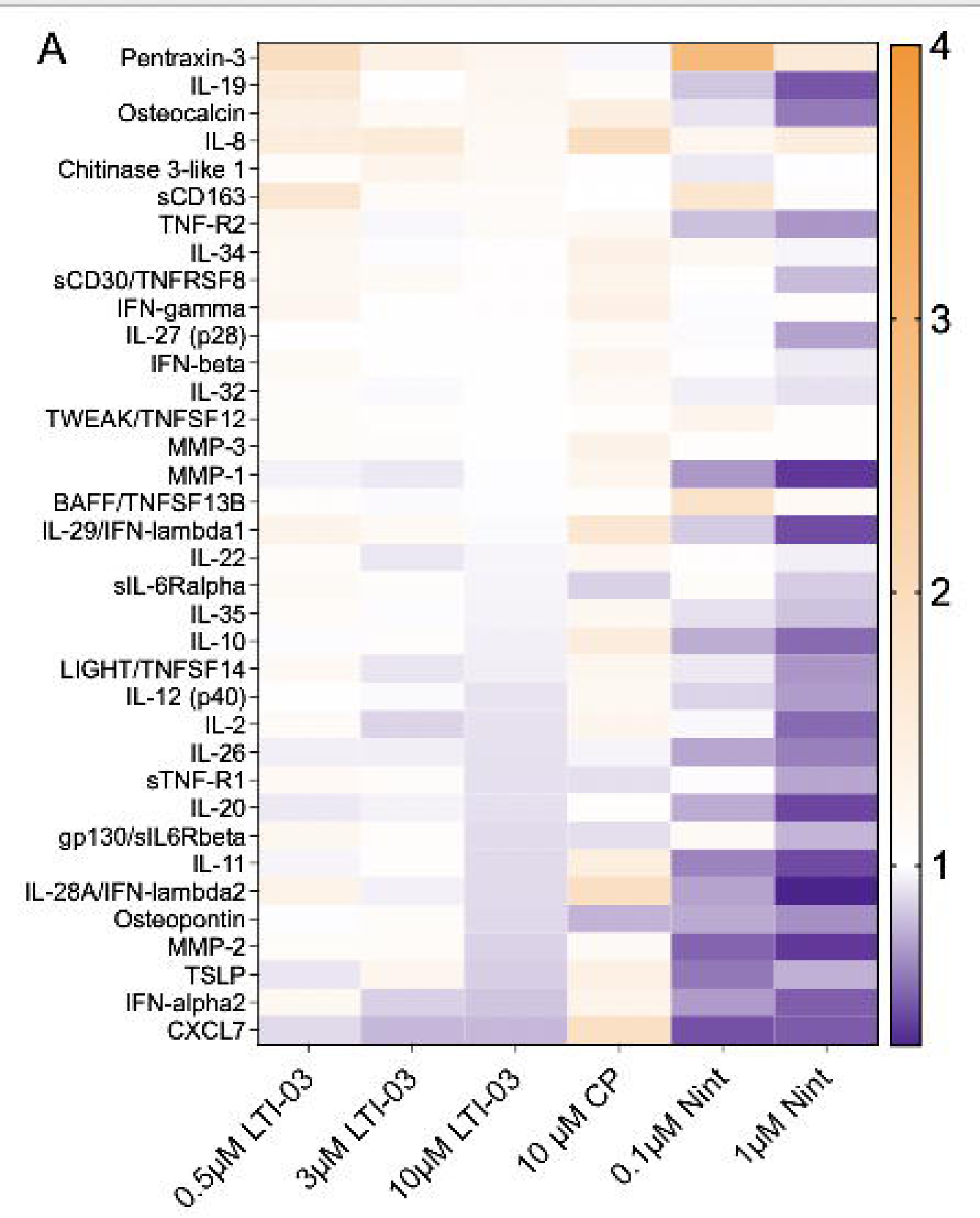
LTI-03 inhibits profibrotic and inflammatory mediators in IPF PCLS following 5 days of treatment. Supernatant was harvested from days 4 or 5 post treatment with 0.5 μM, 3.0 μM, 10.0 μM LTI-03, 10 μM CP or 0.1 μM or 1 μM Nintedanib. (n=8/PCLS per treatment group, except 0.1uM Nintedanib; n=4 PCLS). Heatmap indicates the mean values of per protein measured normalized to the mean of untreated controls.

**Figure 4.**
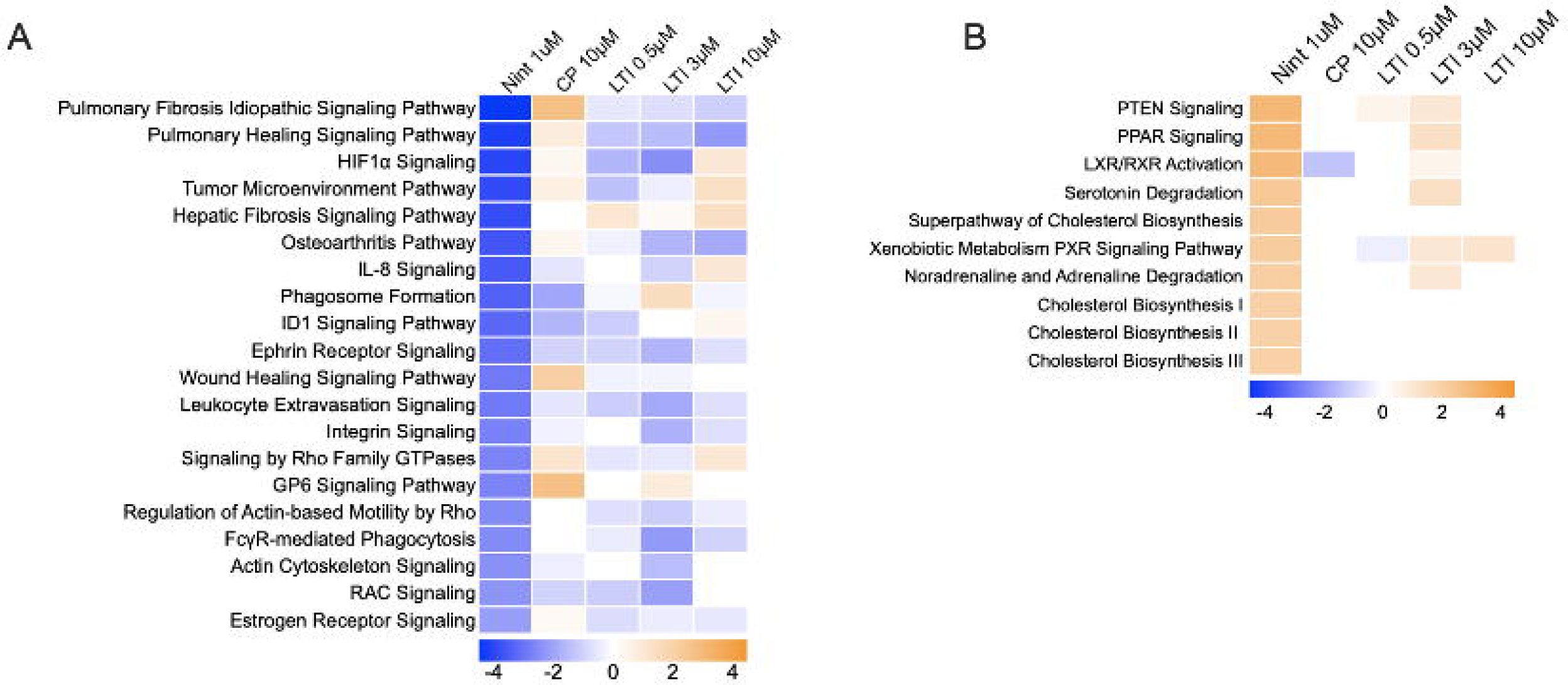
Canonical pathways modulated by nintedanib or LTI-03 treatment of cultured PCLS. (A) Treatment inhibited canonical pathways related to fibrosis signaling in IPF PCLS tissue followed by 5 days of Nintedanib (1.0μM), control peptide (10.0 μM) or LTI-03 (0.5 μM, 3.0 μM, 10.0 μM) treatment every 12 hours. (n=4/group) (B) Treatment activated canonical pathways related to fibrosis signaling in IPF PCLS tissue followed by 5 days of nintedanib (1.0 μM), control peptide (10.0 μM) or LTI-03 (0.5 μM, 3.0 μM, 10.0 μM) treatment every 12 hours. (n=4/group) All values were normalized to untreated groups.

### Expression of upstream fibrosis signaling regulators in cultured IPF PCLS are modulated by nintedanib or LTI-03 treatments

IPA of RNAseq datasets identified key upstream transcriptional regulators related to fibrosis in IPF PCLS cultures after 5 days of nintedanib (1.0 μM), control peptide (10.0 μM) or LTI-03 (0.5 μM, 3.0 μM, 10.0 μM) treatment (n=4/group) (**Figure 5**). All the upstream regulators were inhibited by nintedanib or LTI-03 treatments with the exceptions of insulin growth factor-1 (IGF1), MYC, and KRAS in the LTI-03 treatment groups. Similarly to canonical pathway regulation, 10 µM LTI-03 treatment effects were slightly attenuated as compared to 3 µM LTI-03. All values were normalized to untreated groups. Day 2 analysis is available in **Supplementary Figure 3**.

**Figure 5.**
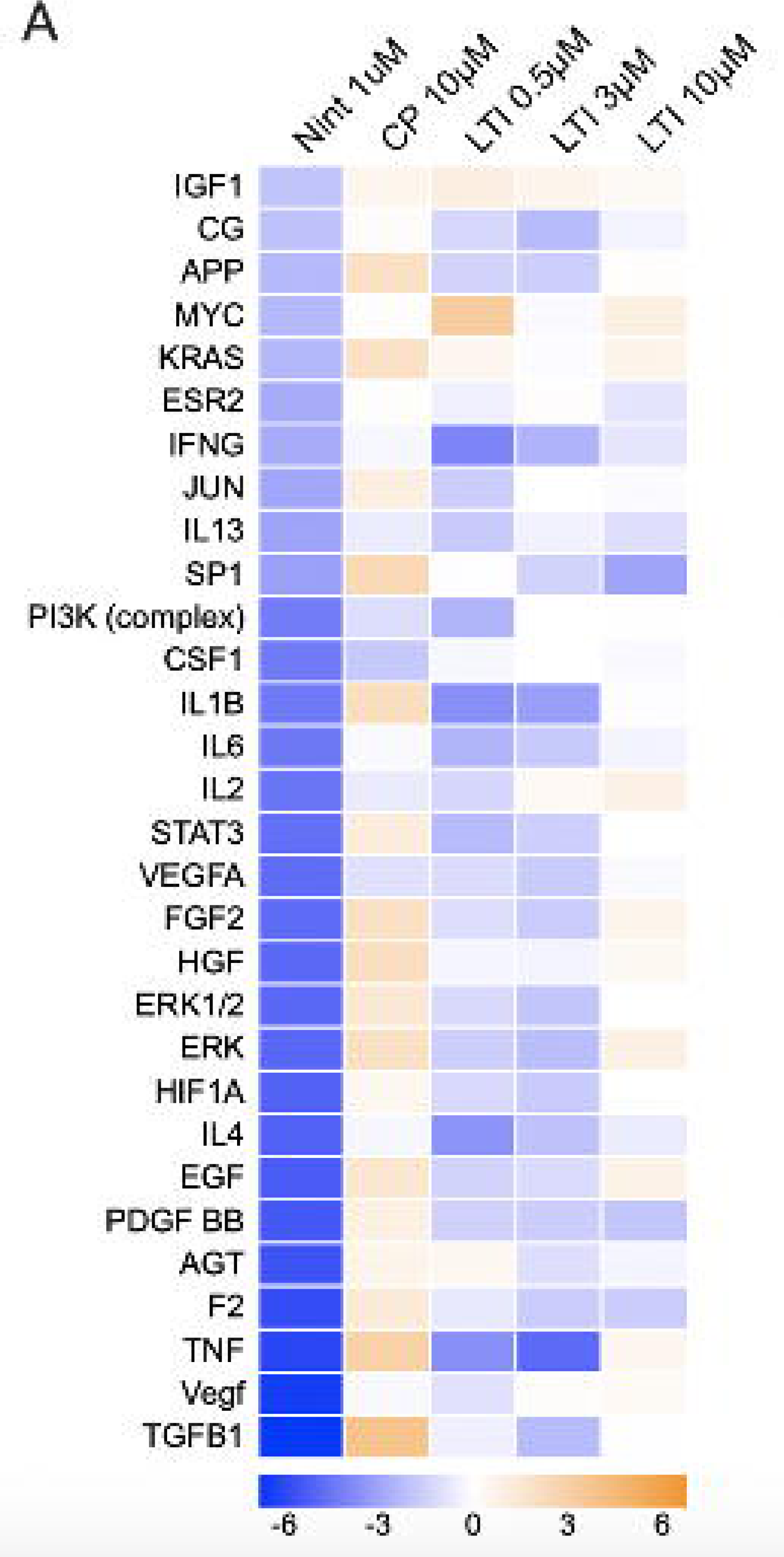
Expression of upstream regulators related to fibrotic signaling in IPF PCLS treated with control peptide, Nintedanib or LTI-03. (A) Expression of upstream regulators genes related to fibrosis pathway in IPF PCLS tissue followed by 5 days of nintedanib (1.0 μM), control peptide (10.0 μM) or LTI-03 (0.5 μM, 3.0 μM, 10.0 μM) treatment every 12 hours. (n=4/group). All values were normalized to untreated groups.

### Modulated expression of disease pathways expression related to fibrotic signaling in IPF PCLS cultures treated with Nintedanib or LTI-03

IPA of RNAseq datasets revealed that disease pathway scores were altered in IPF PCLS cultures after Nintedanib (1.0 μM), control peptide (10.0 μM), or LTI-03 (0.5 μM, 3.0 μM, 10.0 μM) treatment for 5 days (n=4/group) (**Figure 6**). Although there were discrepancies between the 3 and the 10µM dose of LTI-03, Nintedanib and CP increased necrosis and apoptosis pathways, whereas 0.5 and 10 µM LTI-03 suppressed them. Vice versa, cellular homeostasis pathways were rather found to be increased or unchanged in 0.5 or 10 µM LTI-03, but suppressed in nintedanib and 3 µM LTI-03 treated PCLS (**Figure 6**). In addition, both nintedanib and LTI-03 at 3 µM decreased transcript expression associated with Cell proliferation of tumor cell lines, Invasion of tumor cell lines, Invasion of cells, and Cell Movement. Production, Metabolism, & Synthesis of Reactive Oxygen Species was attenuated only by nintedanib treatment. All values were normalized to untreated groups. Day 2 analysis is available in **Supplementary Figure 4**.

**Figure 6.**
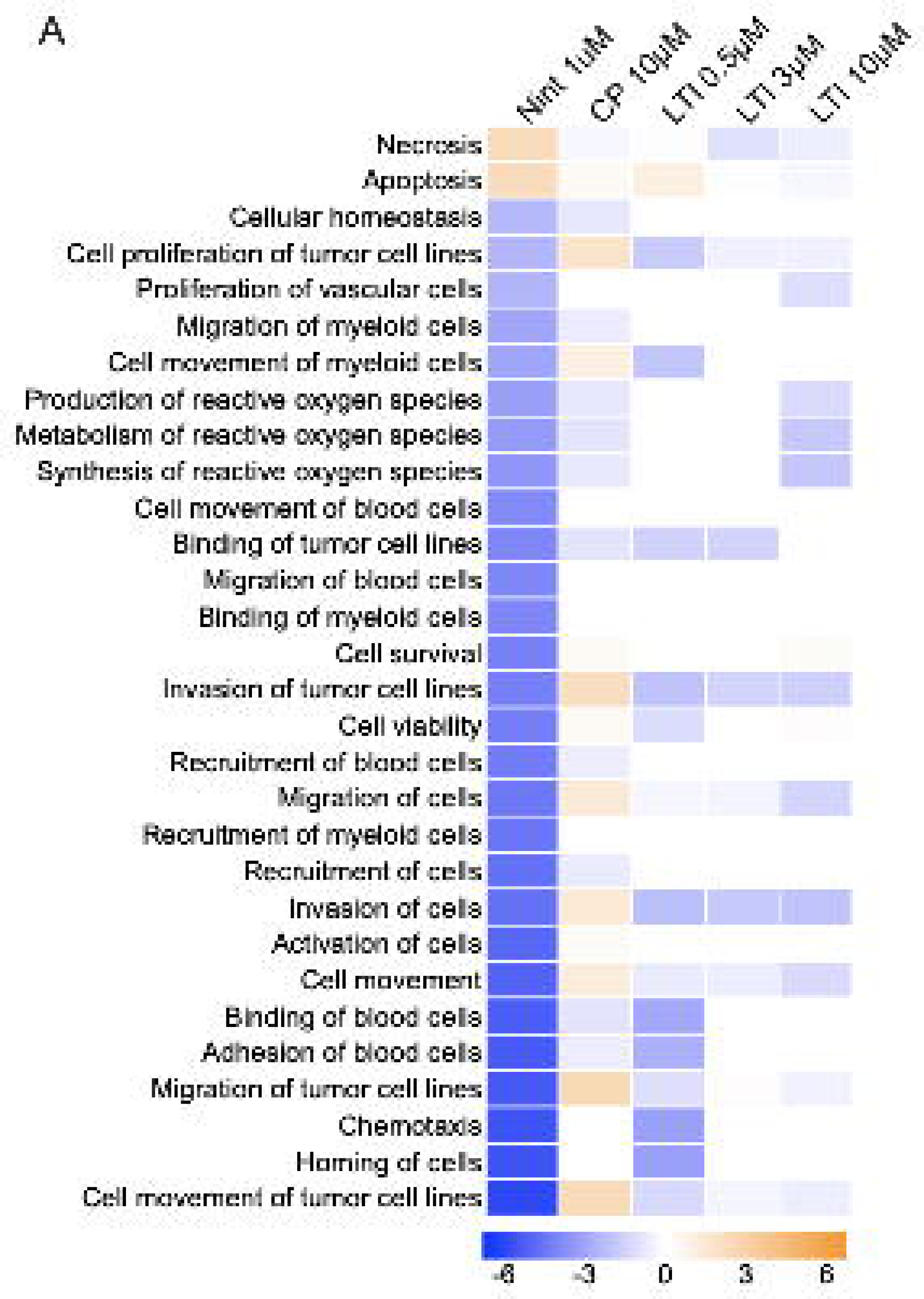
Disease pathways expression related to fibrotic signaling in IPF PCLS treated with control peptide, Nintedanib or LTI-03. Disease pathway expression related to fibrotic signaling in IPF PCLS tissue followed by 5 days of Nintedanib (1.0 μM), control peptide (10.0 μM) or LTI-03 (0.5 μM, 3.0 μM, 10.0 μM) treatment every 12 hours. (n=4/group). All values were normalized to untreated groups.

## Discussion

While countless anti-fibrotic agents are efficacious in animal models of lung fibrosis, most putative therapeutic interventions continue to fail in IPF clinical trials (25). Although there are several interventional therapies currently in clinical development that are narrowly focused on pro-fibrotic or lung remodeling pathways, (30)(31) to date only drugs such as nintedanib (27) and pirfenidone (28) which demonstrate inhibitory effects on multiple pathways have successfully slowed the rate of lung function decline in IPF patients. Consequently, experimental models are required that are more predictive of clinical success. The culture of PCLS generated from explanted IPF patient lungs has emerged as a promising *ex vivo* translational approach to study drug effects directly on IPF tissue. (26) As compared to animal models of lung fibrosis, such approach may offer distinct advantages: 1) IPF PCLS are derived from humans, 2) the diseased tissue contains maintains the same perpetuating fibrotic mechanisms driving human IPF onset and progression, 3) they provide a translational platform in which to study all the cell types present in the IPF lung as well as the extracellular matrix environment that encompasses these cell types (43), 4) they represent the complex genetic/epigenetic background of IPF patients, 5) they originate from IPF patients with a defined exposome history, and 6) placebo / vehicle treated PCLS can be included as experimental controls for each experiment and each patient. As a result, the IPF PCLS system may be much more informative with respect to the impact of putative therapeutic agents in human IPF (44–47).

In the present study, the IPF PCLS system was characterized by dynamic temporal changes in both pro-fibrotic proteins and transcripts, reflecting the progressive nature of the disease. Many of the highly upregulated soluble protein factors in IPF PCLS cultures have been described in lung fibrosis including thymic stromal lymphopoietin (TSLP) (48), osteopontin (OPN) and chemokine CXCL7 (49). OPN is known to bind CD44 and trigger the phosphorylation of FAK, a profibrotic pathway in the PCLS assay. (51) Transcriptional analysis via bulk RNA sequencing of IPF PCLS also showed that profibrotic intracellular/plasma membrane, ECM, and secreted factors were upregulated in a temporal manner in the IPF PCLS system, again reflecting perpetuation of fibrosis. Many of the dynamic changes in transcript expression were tied to ECM components, most notably collagens. Thus, the PCLS culture system provides a clinically relevant approach to study pro-fibrotic mechanisms and targeting strategies directed at these mechanisms in IPF.

To gain insight into the mechanism of action and putative clinical efficacy of LTI-03, the IPF PCLS system was used to evaluate the antifibrotic activity of physiologically relevant doses of LTI-03 compared to the standard of care drug nintedanib. Due to epigenetic silencing (29), loss of transcriptional regulators such as FOXO3A (30), and other mechanisms (5), CAV-1 expression is markedly reduced in IPF (29, 31) and CAV-1 is a prognostic predictor in IPF (32). While a gene transfer technique has been used to restore Cav-1 protein expression and ameliorate fibrosis in an experimental lung fibrosis model (31), the antifibrotic effects of CSD peptides have been demonstrated in pulmonary fibrosis models (16, 33, 34) as well as fibrosis due to aging in mice (35). CSD peptides have modulatory effects on monocultures of primary IPF fibroblasts (16) and epithelial cells (unpublished), but the effect of these peptides in primary multi-cellular systems has not been explored to date. We observed that LTI-03 treatment dose-dependently reduced the deposition of COL1A1 protein in IPF PCLS at day 5. While nintedanib has been shown to reduce COL1A1 transcript expression in normal lung PCLS exposed to a fibrotic cocktail (36) and modulates neoepitope biomarkers of type III collagen turnover (37), we did not observe a statistically significant change in COL1A1 protein in the IPF PCLS system after 5 days of treatment, again lending credit to the concept that human IPF PCLS may more appropriately reflect the human IPF disease as compared to PCLS derived from a rather healthy lung treated with a profibrotic cocktail.

LTI-03 has the potential to modulate several canonical intracellular signaling pathways due to its putative interaction with approximately 30% of all proteins; those that contain CBDs (38, 39). The seven amino acid sequence of LTI-03 is naturally occurring as it is derived from the CSD portion of CAV-1, which is comprised of amino acid sequences 82-101 of the NH_2_ terminal region. This region plays a role in CAV-1 dimerization as well as regulation of diverse signaling intermediates, many of which are implicated in the pathogenesis of fibrosis (4, 40, 41). In addition, caveolae work to regulate endocytosis as well as many receptors signaling pathways. In healthy lung cells, endogenous caveolin-1 modulates receptor signaling towards homeostasis via facilitation of receptor stabilization, turnover and signaling (42). Homeostatic modulation is conferred via interactions between CSD of caveolin-1 and the CBD of many receptors and proteins in the plasma membrane. During fibrosis, increased growth factor signaling disrupts cellular homeostasis leading to loss of endogenous CAV-1 expression and destruction of caveolae. Consequently, we hypothesize that the loss of CAV-1 as a homeostatic signaling mediator in the IPF lung is a major contributor to aberrant signaling present in IPF patients. Our *in-silico* analysis of endogenous proteins containing CBDs highlighted many growth factor receptors including FGFR2, VEGF, PDFRA/B, among others, which are relevant to IPF pathogenesis (24). LTI-03 likely confers broad attenuation of profibrotic signaling, in part, via its regulatory effects on aberrantly activated growth factor receptors (**Figure 7**). While direct binding interactions of LTI-03 with these endogenous factors have not been studied to date, herein we provide both protein and transcript analyses suggesting that LTI-03 is targeting multiple receptor-directed pathways in IPF PCLS cultures. Also, overexpression of Cav-1 had been shown before to improve barrier function and reduce TSLP expression in cultured human airway epithelial cells. (50)

**Figure 7.**
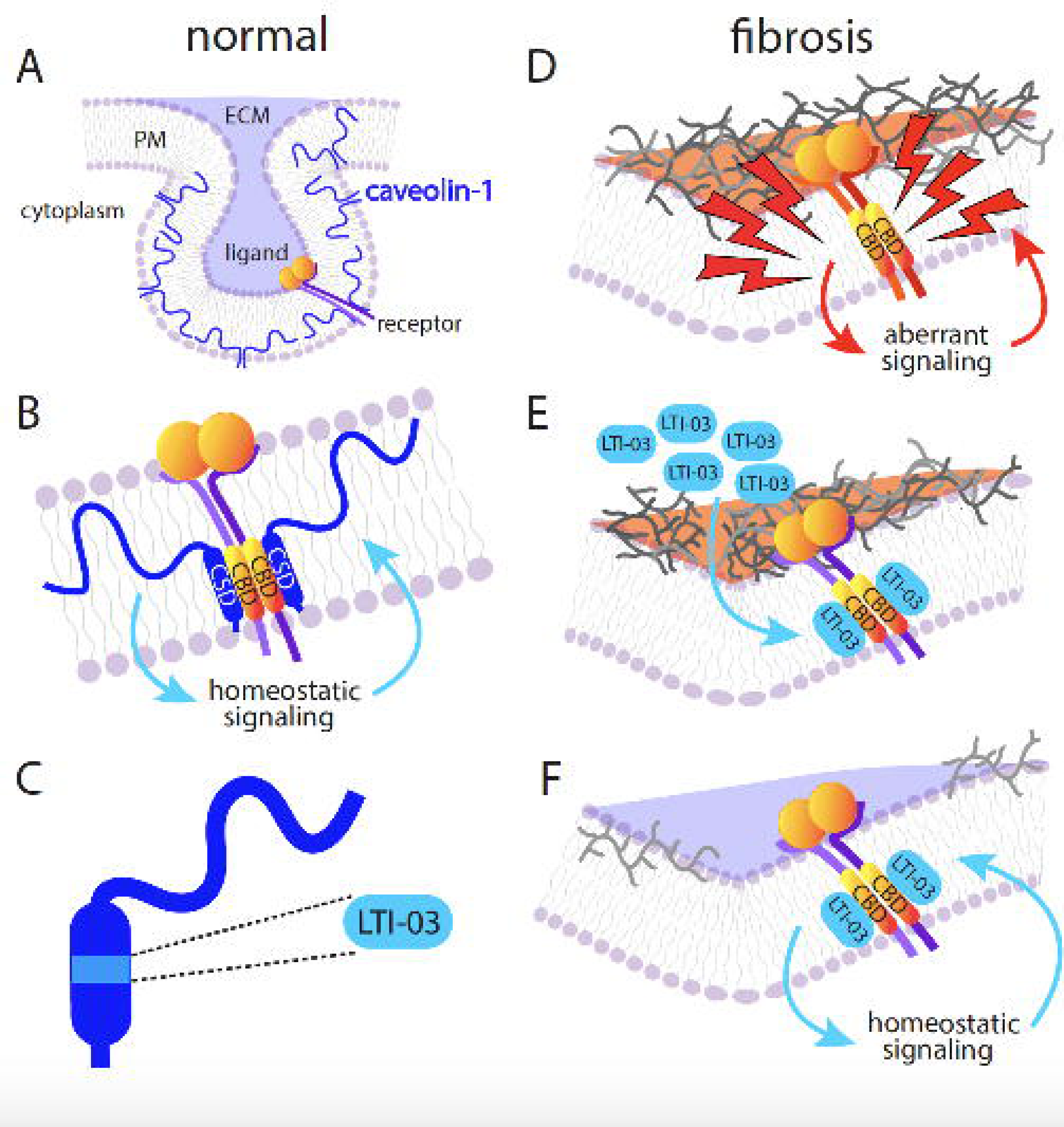
Putative mechanism of LTI-03 anti-fibrotic activity in end stage IPF PCLS. (*A*) In normal lung cells, caveolin-1 (CAV1) is robustly expressed in AEC2s, AEC1s, bronchio-epithelium and lung endothelium Caveolin-1 is positioned in the plasma membrane (PM) where it supports the formation of PM invaginations known as caveolae that are thought to be important for mechanical stretching of the lung and which function as receptor sinks. (*B*) The caveolin scaffolding domain (CSD) is comprised of amino acid sequences 82-101 of the NH_2_ terminal region and plays a role in CAV-1 dimerization as well as regulation of diverse signaling intermediates, many of which are implicated in the pathogenesis of fibrosis. The CSD domain is known to interact with receptors and proteins that contain a caveolin binding domain (CBD) and interaction can result in stabilization, internalization, or recycling of the membrane proteins thereby contributing to homeostatic signaling. (*C*) LTI-03, the 7-mer peptide used in this study and currently in clinical trials for IPF, is derived from the CSD portion of caveolin-1 (amino acid sequences 89 to 95). (*D*) After initiation of progressive lung disease (indicated by lightning bolts), extracellular matrix proteins are deposited which perturb lung architecture and generate aberrant signaling and gene expression. Caveolin protein expression in fibrotic lungs are significantly decreased which likely contributes to aberrant cell signaling. (*E*) In the context of lung fibrosis, LTI-03 may act as a surrogate CSD which results in abrogation of aberrant signaling of local receptors regardless of cell type. The putative interaction between LTI-03 and aberrantly profibrotic receptors or signaling proteins containing CBDs leads to a reduction of profibrotic gene expression and mediators which results in a decrease in extracellular matrix deposition and return to cellular homeostasis.

Stegmayr and colleagues have highlighted the challenges associated with obtaining high-yield and -quality RNA for RNA-sequencing of human PCLS (52). In this study, PCLS was generated from 4 IPF patients and high quality RNA was isolated from untreated and treated PCLS. While nintedanib was more potent than LTI-03 in the modulation of various fibrosis-related canonical pathways and upstream regulators, LTI-03 exhibited a dose-dependent breadth of modulation that overlapped considerably with the SOC drug. However, LTI-03 did not promote cellular necrosis or apoptosis that were observed in nintedanib-treated PCLS cultures. Thus, the present study confirms the utility of the IPF PCLS system for RNA sequencing and analysis to address fibrotic processes and the efficacy of therapeutic approaches in IPF.

Future studies using single cell RNA sequencing would unmask cell-specific transcriptional changes following either LTI-03 or nintedanib treatment in the PCLS system. In addition, a detailed comparison, and characterization of fresh IPF PCLS explant tissue with previously frozen tissue may allow for the accommodation of more robust sample numbers in the future; and would also accommodate the procurement of equal numbers of male and female derived tissues. Finally, we were limited in our analysis of epithelial-associated markers and factors given that the particular IPF lung explants used to generate PCLS used in this study were largely devoid of viable type 1 and 2 epithelial cells. Studies are ongoing to explore the effect of LTI-03 on lung epithelial cell biology in IPF.

## Conclusion

In summary, the expression profile of untreated IPF PCLS over five days in culture suggested dynamic changes in fibrotic pathways rendering this translational model an appropriate model for testing clinically relevant antifibrotic agents. Consistent with earlier studies (53), this study indicates a pivotal role for the use of caveolin-1 scaffolding domain peptides for the therapeutic intervention of lung fibrosis and highlights LTI-03’s potential therapeutic significance for IPF patient. This study is the first to characterize the dose-dependent effect of LTI-03 on critical canonical signaling pathways active in an IPF PCLS system. Finally, LTI-03 exhibited a similar pattern of inhibition and activation compared with nintedanib treatment but without the noted toxicity observed with the multi-tyrosine kinase small molecule inhibitor. LTI-03 was well-tolerated in non-diseased volunteers (NCT04233814) and a Phase 1b safety trial in IPF patients (NCT05954988) is currently enrolling.

## Acknowledgements

This work was funded partially funded by the National Institute of Health (NIH) grant P01-HL108793 and Lung Therapeutics, Inc.

## Supplementary Figures

**Supplementary Figure 1 LTI-03 inhibits profibrotic and inflammatory mediators in IPF PCLS following 2 days of treatment** Supernatant was harvested 2 days post treatment with 0.5 µM, 3.0 µM, 10.0 μM LTI-03, 10 μM CP or 0.1 μM or 1 μM Nintedanib. (n=8/PCLS per treatment group, except 0.1 μM Nintedanib; n=4 PCLS). Heatmap indicates the mean value per protein measured and normalized to the mean of untreated controls.

**Supplementary Figure 2. Expression of canonical pathways related to fibrotic signaling in IPF PCLS treated with control peptide, Nintedanib or LTI-03 after 2 days of treatment.** (A) Activated canonical pathways related to fibrosis signaling in IPF PCLS tissue followed by 5 days of nintedanib (1.0 μM), control peptide (10.0 μM) or LTI-03 (0.5 μM, 3.0 μM, 10.0 μM) treatment every 12 hours. (n=4/group)

**Supplementary Figure 3. Expression of upstream regulators related to fibrotic signaling in IPF PCLS treated with control peptide, Nintedanib or LTI-03.** (A) Expression of upstream regulators signaling related to fibrosis signaling in IPF PCLS tissue followed by 2 days of nintedanib (1.0 μM), control peptide (10.0 μM) or LTI-03 (0.5 μM, 3.0 μM, 10.0 μM) treatment every 12 hours. (n=4/group). All the values are related to DEGs obtained from the comparison between each treatment to untreated groups.

**Supplementary Figure 4. Disease pathways expression related to fibrotic signaling in IPF PCLS treated with control peptide, Nintedanib or LTI-03.** Disease pathway expression related to fibrotic signaling in IPF PCLS tissue followed by 2 days of Nintedanib (1.0 μM), control peptide (10.0 μM) or LTI-03 (0.5 μM, 3.0 μM, 10.0 μM) treatment every 12 hours. (n=4/group). All the values are related to DEGs obtained from the comparison between each treatment to untreated groups.

