## Supplementary figures and images for "Pre-clinical proof-of-concept of anti-fibrotic activity of caveolin-1 scaffolding domain peptide LTI-03 in *ex vivo* precision cut lung slices from patients with Idiopathic Pulmonary Fibrosis"

### Supplemental Figure 1

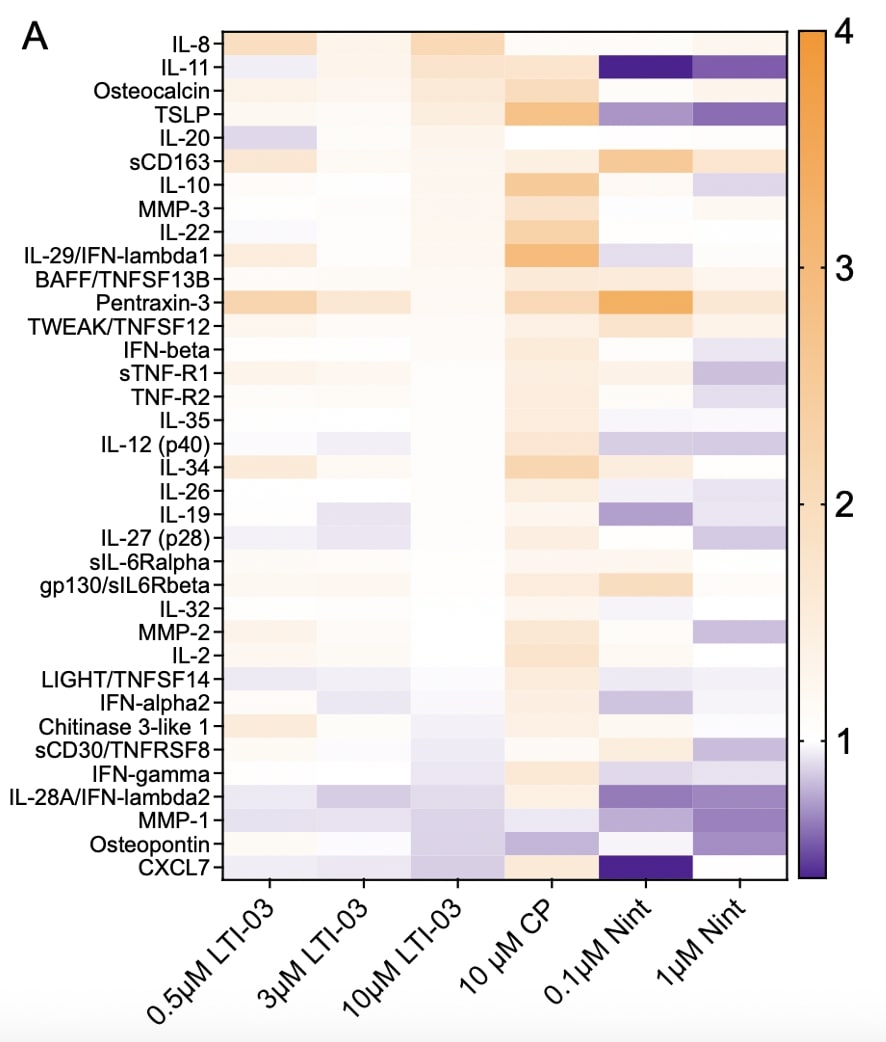

### Supplemental Figure 2

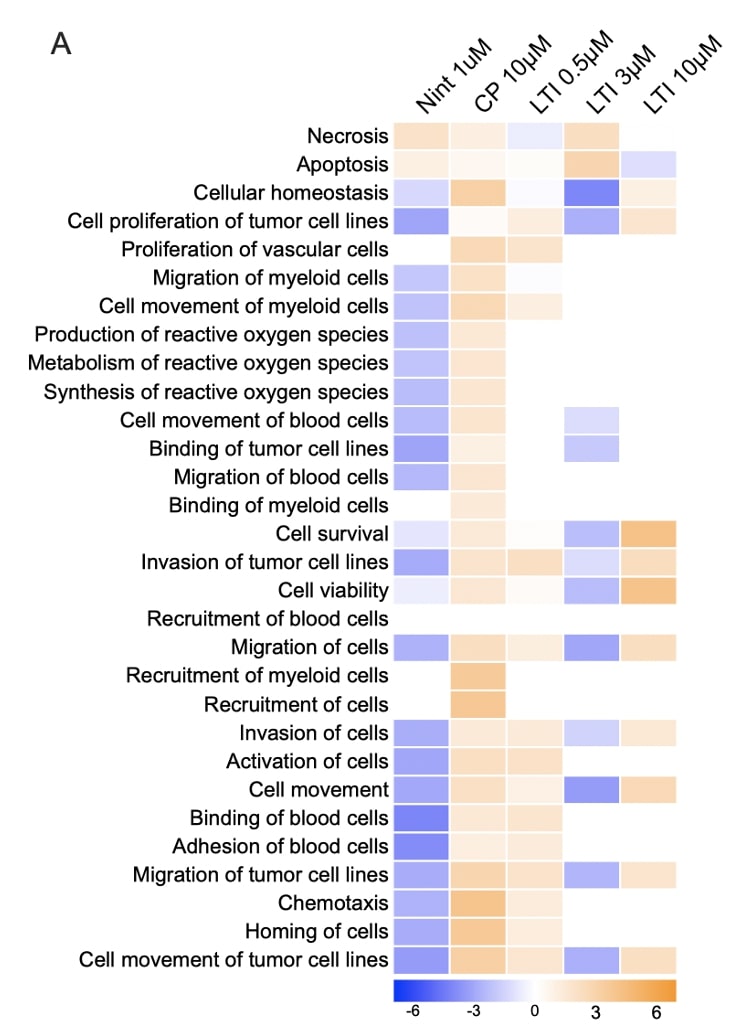

### Supplemental Figure 3

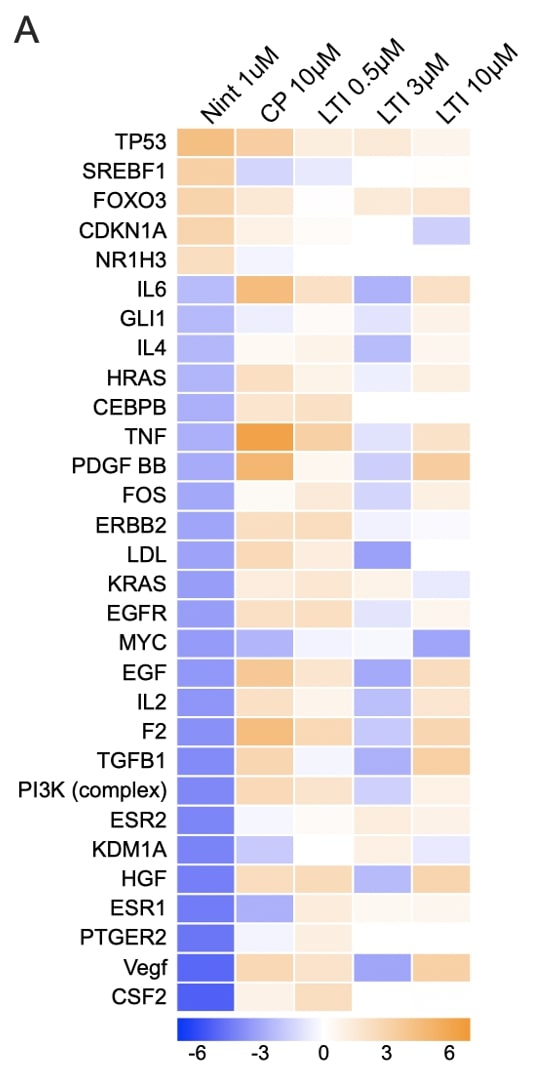

### Supplemental Figure 4

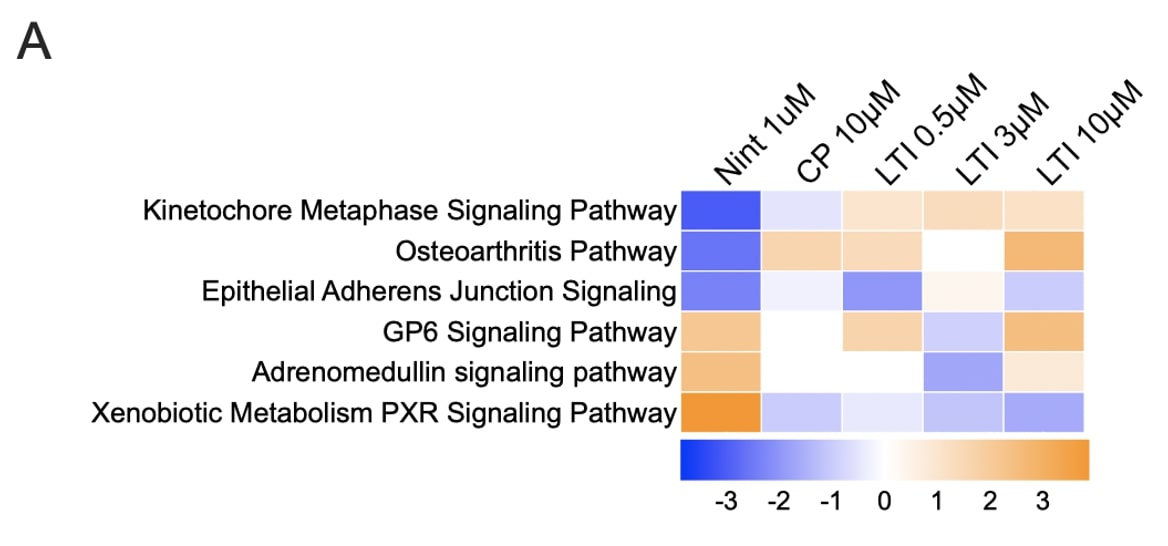
